# Predicting resprouting of *Platanus* × *hispanica* following branch pruning by means of machine learning

**DOI:** 10.1101/2023.08.11.552927

**Authors:** Qiguan Shu, Hadi Yazdi, Thomas Rötzer, Ferdinand Ludwig

## Abstract

- Resprouting is a crucial survival strategy following the loss of branches, being it by natural events or artificially by pruning. The prediction of resprouting patterns on a physiological basis is a highly complex approach. However, trained gardeners try to predict a tree’s resprouting after pruning purely based on their empirical knowledge and a visual check of the tree’s geometry. In this study, we explore in how far such predictions can also be made by algorithms, especially using machine learning.
- Table-topped annually pruned *Platanus* × *hispanica* trees at a nursery were documented with terrestrial LiDAR scanners in two consecutive years. Topological structures for these trees were abstracted from point clouds by cylinder fitting. Then, new shoots and trimmed branches were labelled on corresponding cylinders. Binary and multiclass classification models were tested for predicting the location and number of new sprouts.
- The accuracy for predicting whether having or not new shoots on each cylinder reaches 90.8% with the LGBMClassifier, the balanced accuracy is 80.3%. The accuracy for predicting the exact numbers of new shoots with GaussianNB model is 82.1% but its balanced accuracy is reduced to 42.9%.
- The results were validated with a separate evaluation dataset. It proves a feasibility in predicting resprouting patterns after pruning using this approach. Different tree species, tree forms, and other variables should be addressed in further research.

## 1. Introduction

Disturbances to tree growth like ice storms, fires, wind and diseases (Hauer et al., 2006; Simler et al., 2018) are common in nature. They cause great loss in trees’ biomass especially above the ground. In view of this, resprouting is a vital survival strategy for most tree species: new shoots can grow out of dormant buds rapidly at certain positions after the disturbance occurred. This process is recognized as a major force in forest regeneration (Matula et al., 2019) and has significant impacts on forest dynamics (Martini et al., 2008). Humans recognized and harnessed these phenomena from early times (Candel-Pérez et al., 2022; Petit & Watkins, 2003). A famous example is pollarding, where all the shoots of a tree crown are regularly cut off to encourage the growth of new sprouts, which were used as firewood and material for weaving baskets.

Regardless of the practical use, it is a highly interesting but at the same time very complex challenge to understand and predict the resprouting patterns of trees caused by disturbances on a physiological basis. These patterns are firstly determined by axillary buds which either form new shoots or enter dormancy (Suzuki, 2002). This “decision” is essentially controlled by hormone signals. Auxin was considered one of the main mediator in the 20th century, while new findings indicate that cytokinins (Salam et al., 2021; Schneider et al., 2022) and strigolactones (Gomez-Roldan et al., 2008) play a major role in apical dominance and branching inhabitation respectively. Without a clear conclusion yet regarding their exact mechanisms, studies tried to understand resprouting patterns from other micro and macro perspectives: its relation to genetic regulation (Hill & Hollender, 2019), in responding to seasonal adaptation (Singh et al., 2022), or by an explanation known as Low Energy Syndrome (Martín-Fontecha et al., 2018).

However, these endogenous physiological processes do not tell the whole story of resprouting. Leaf area and light are redistributed after the disturbances, which then affects photosynthetic processes (Balandier et al., 2000). This does not simply mean a decrease in photosynthetic capacities, but involves reallocation of carbon- and other resources among plant organs such as fruits (Kohek et al., 2015; Tosto et al., 2023) and flowers (Grecia et al., 2022). What makes the impact of this disturbance even more complex is timing. For example, summer pruning on an apple tree normally causes a temporary loss of apical dominance and increase its cytokinin supply. But depending on its exact timing, the dominance may be delayed or even prevented (Saure, 1987). As a result, a precise analysis of how a disturbance reshapes a tree using a physiological approach must address 1) primary status of hormone, 2) resource reallocation, and 3) timing issue. To our knowledge no research has brought all these aspects together so far.

Even without any precise analytical tools regarding resprouting analysis, skilled practitioners learn how to prune a tree in their charge. They neither measure its sap-flows with multiple sensors nor meter the cytokinin concentration in chemistry labs. By going around the tree and observing the main branches, they decide where to prune. Their decisions are based on empirical knowledge of natural phenomena, originally derived from accurate observations of causes and effects – the tree’s resprouting reaction to the loss of branches by pruning. Countless repetitions of similar processes have been experimented in horticulture over centuries (Saunders, 1898). For a gardener, his or her primary pruning skills may start with a set of general rules written in a manual book (Brickell & Joyce, 1996). Then, their skills will independently evolve further through repeated practices at work specific to different climate zones, species etc. If their pruning decisions lead to resprouting reactions largely similar to their expectations, gardeners finally prove to be able to predict the tree’s reaction purely on visual observation and geometrical patterns without digging deep into simulating physiological processes.

In horticulture and arboriculture, we currently see a strong trend towards automation of pruning by machines or robots (Sam et al., 2022). So far, these are comparatively simple, standardized actions (M. Li et al., 2021; Sam et al., 2022), but the more complex the tasks become in this regard, the more important is a plausible, robust and prompt prediction of the growth reaction of a tree to pruning. At the same time, it can be assumed that in the future trees worldwide will increasingly experience growth disturbances due to the consequences of climate change (drought, stronger and more frequent storms), which will be coupled with a loss of biomass and subsequent resprouting. In order to assess the development of such trees, for example in an urban context, also here a plausible, robust and prompt prediction of resprouting in response to the previous loss of branches and twigs is necessary.

In this regard, physiological forecasts are too complex, rely on too many often-unknown parameters (e.g. weather) and thus are too sensitive to errors and too slow. The aim of the study at hand is to develop first basics for a prediction model on the basis of geometric patterns corresponding to the approach of experienced gardeners using a concrete example.

Rapid development in remote sensing is providing a solid base for this aim. First of all, terrestrial LiDAR scanners can capture detailed geometry of objects with a precision up to 3 mm from multiple standing positions (RIEGL, 2023). This method proves capable of capturing a tree’s trunk and branches with more than 10 mm diameter (Gobeawan et al., 2018; Yang et al., 2022) during its leaf-off state (Kükenbrink et al., 2022). Raw data is stored in the form of a discrete point cloud. Furthermore, different approaches have been developed to extract tree structure: skeleton abstraction following occupancy grids (Bucksch et al., 2010; Sun et al., 2022); branch direction by eigenvectors of point patches or sections (Bremer et al., 2013; Raumonen et al., 2013); skeleton as the Dijkstra’s shortest path from the tree base to ends (Du et al., 2019; J. Li et al., 2022, 2022; Wang et al., 2014); skeleton redrawn with searching steps (Hackenberg et al., 2014); learning the reconstruction pattern through a neural network (Liu et al., 2021). Overall, these abstracted information about tree architecture is called quantitative structure model (QSM) (Åkerblom et al., 2017; Shu et al., 2022). In this way, every segment of the tree stem or branch can be retrieved containing its diameter, length, axial direction, hierarchy in the whole branching structure as well as the pointer to its parent and child segments.

These data for a computational model are like experiences for a human brain. The process for an algorithm to “learn from experience” without being explicitly programmed was defined as machine learning (Samuel, 1959). Over 70 years of development, machine learning models have proven capable and efficient to inherently solve the 5 typical problems of data science, namely classification, anomaly detection, regression, clustering and reinforcement learning (Alzubi et al., 2018). Among them, classification models can assign class labels to testing instances where the predictor features are known (Kotsiantis, 2007). Depending on the number of output labels, they work for both binary and multi-class classification problems. To establish such models from a practical aspect, open-source packages such as scikit-learn (Pedregosa et al., 2011) have integrated common ones, offering an easy access to adapt parameters for different applications.

Equipped with the digital tools above, accurate information regarding tree structures can be collected and processed in analogy to what a real gardener does. Building on this, we are addressing the following questions: How can we predict the position and number of resprouting shoots based on a purely “visual approach” (pattern recognition)? Which machine learning model achieves the best accuracy for this task?

## 2. Materials and Methods

### 2.1 Study case

To address our questions, we looked for tree cases that are frequently pruned in a distinct manner under similar environmental conditions. At Bruns Nursery, Bad Zwischenahn in the north of Germany, so-called table-topped plane trees (*Platanus × hispanica*) are grown in a clearly defined area under standardized conditions. The crowns of these tree are shaped into a flat layer through labour-intensive maintenance. This form probably origins from Baroque gardens, where plants were kept in an orthogonal manner to enhance the orientation or perspective (Dobrilovič, 2010). Due to the expansion of the crown like an umbrella, it is still used in European cities nowadays for shading squares and pedestrian areas (e.g., the central square at Labouheyre, France). To produce such trees, there are in general two phases. In the first phase, a young plane tree with a naturally grown canopy is intensively trimmed. At around 3 meters height six branches are selected and bent horizontally into different directions with equal angles in between. Where necessary, bamboo sticks are added as temporary supports to force the branch into the aimed direction (see “1^st^ year” in Fig. 1). In the second phase, new shoots or even some of the older shoots from these six main branches are carefully selected and pruned by experienced gardeners. Pruning decisions are important at this phase to enable shoot growth only at desired positions. Some shoots reserved from previous years could still be trimmed off if there appears another new shoot that becomes a better option. This procedure is repeated in the following years (see “2^nd^-6^th^ year” in Fig. 1). Multiple reiterations of the tree by resprouting result in a complex branching pattern. Due to the annual pruning and relative complex branching pattern the second phase of these cases is considered effective to analyse abilities of machine learning models in predicting resprouting patterns based on quantitative structural tree models under complex yet repetitive conditions. It should be noticed that the aim of this study is not recreating this specific form of tree geometry like the table-topped *Platanus × hispanica* but to gain fundamental knowledge regarding resprouting reactions of trees.

**Figure 1.**
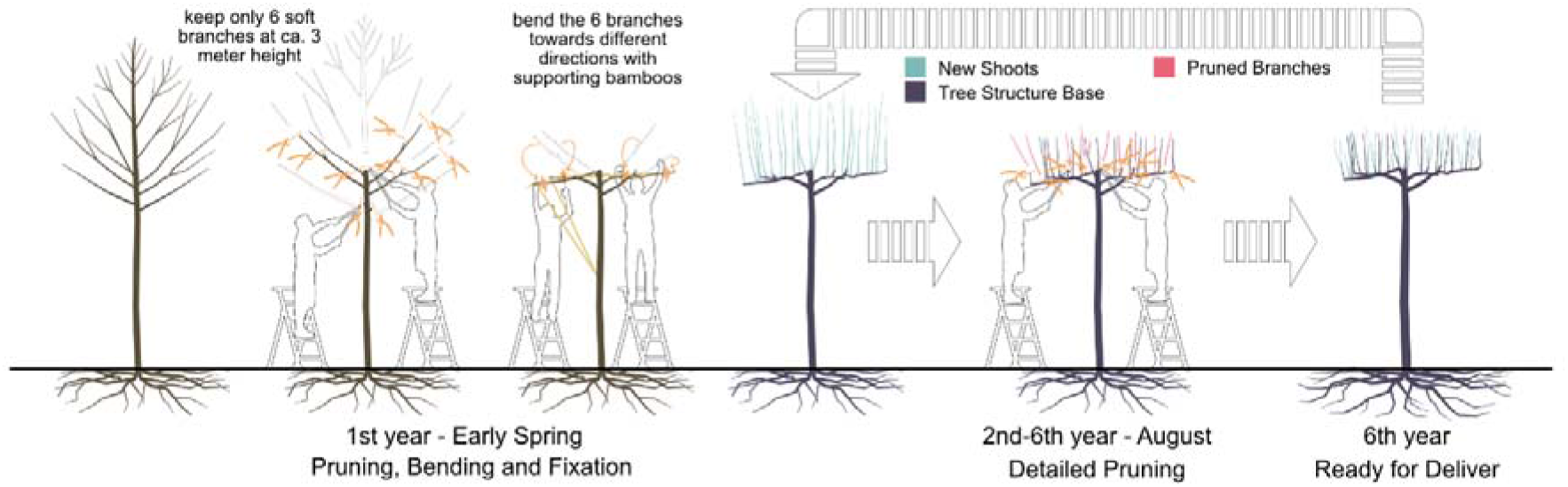
The procedure for producing a table-topped *platanus* through iterative branch and shoot selection and pruning with intensive labour force.

### 2.2 Data acquisition and pre-processing

In subsequent two winters, namely in January 2022 and January 2023, an area consisting of 3- and 4-year-old table topped platanus (see Fig. 2a) planted in 3 rows at Bruns Nursery were scanned with LiDAR scanner RIEGL VZ-400i. The scanner was mounted on a tripod in 2022, while mounted on a vehicle (see Fig. 2b) in 2023. All the scans were set to “Panorama30” standard (with angular resolution 0.030°) and conducted in a “stop-and-go” method. Scanning positions were located along each row at every third tree (ca. 12 m). Point clouds from different scan positions were automatically registered in RiSCAN Pro in reference to GNSS coordinates recorded with Leica Zeno FLX100 plus (Leica, 2023). The original GNSS coordinates indicate accuracies ranging between 0.68 to 0.80 m at different scan positions. Therefore, the reliability on GNSS was set to low during the automatic registration and the multistation adjustment. With all steps above, we got two point clouds containing all the tree cases for the year 2022 and 2023 respectively. Afterwards, individual trees were segmented manually (see Fig. 2c). This manual step is efficient for our cases because those trees planted in the nursery were almost perfectly aligned in an equal distance and their crowns did not touch each other. The ground surface was flat and clear and there were no irrelevant objects such as shrubs around tree trunks. A total of 49 plane trees were scanned in 2022 while the number of trees scanned in 2023 was 28, (due to tree sales during 2022, see Fig. 2d and 2e). As a result, we got point clouds of 28 plane trees for both years.

**Figure 2.**
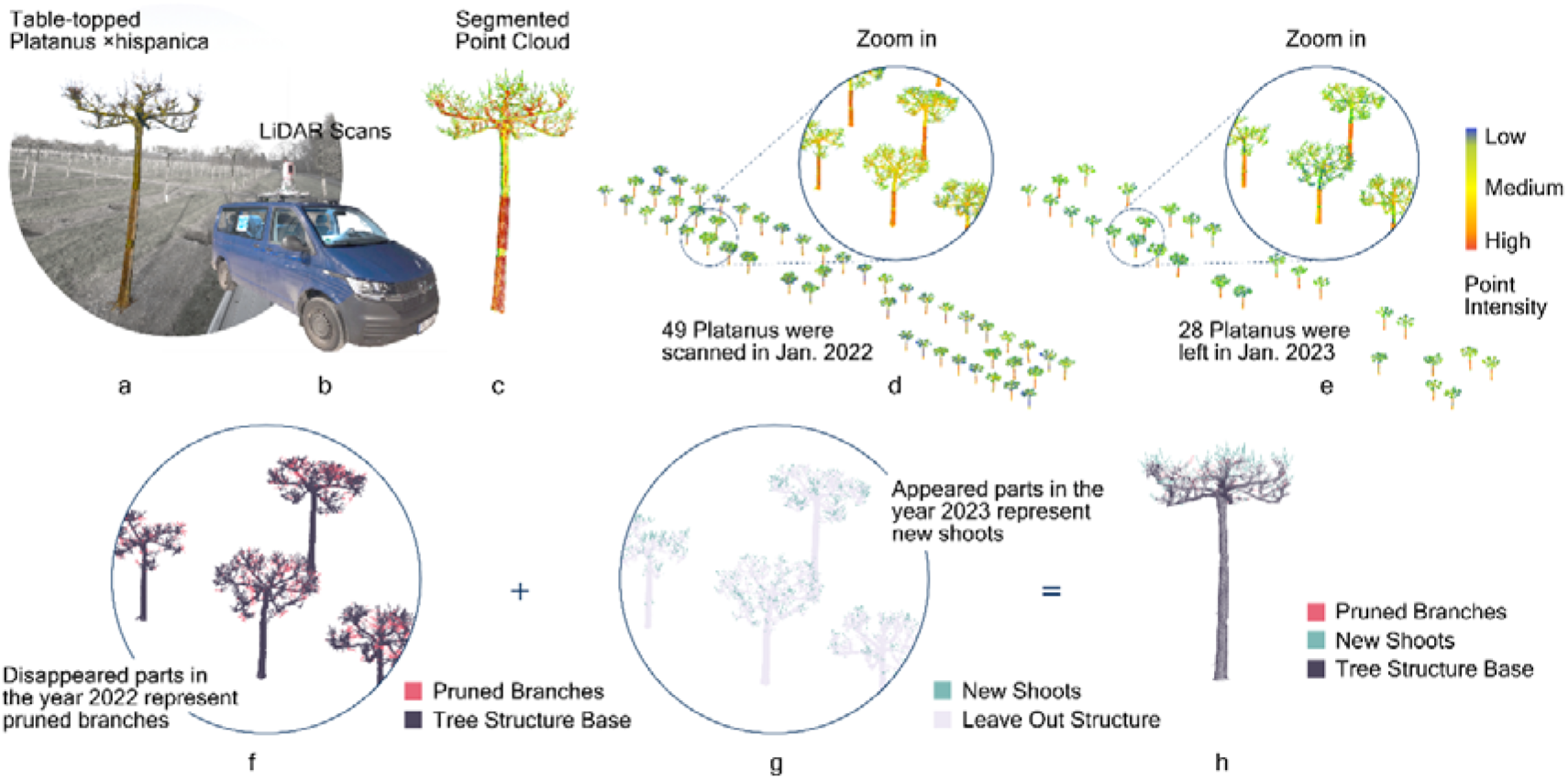
The overall procedure for detecting pruned branches and new shoots from point clouds of LiDAR scans of two consecutive years.

The next step was to identify changes in the geometrical structure of the trees in these two years (see Fig. 2h). For this purpose, the two corresponding scans regarding the same trees must be aligned. The GNSS coordinates have an offset up to 0.8 meters, which is not sufficient to our demand. The most common algorithm for matching 3d models precisely, namely Iterative Closest Point (Rusinkiewicz & Levoy, 2001), does not work for these tree cases because the new shoots and the extensive pruning on tree branches have altered their geometries significantly. A supervised alignment by manually picking point pairs on corresponding branch surfaces also caused visible deviations owing to the girth growth. Finally, we manually aligned all tree pairs individually using multiple views. This guaranteed the best possible alignment despite significant geometrical changes between the two scans. Only then, we were able to precisely detect the changes caused by growth and pruning between the point clouds. In principle, point sets that only appeared in the scan of 2022 and disappeared in the scan of 2023 should represent branches pruned away. Conversely, point sets that were only found in 2023 should represent new shoots. In practice, however, an object even with no change in its geometry is unlikely to have identical points on its surface in the two independent scans. To solve this, cloud-to-cloud distance (Jafari et al., 2017) was applied. It calculates the distance between each point in one point cloud to its nearest neighbour in the other point cloud using Hausdorff distance algorithm (Girardeau-Montaut, 2023). Depending on point cloud quality and precision in alignment, a minimum distance threshold ranging between 0.020 to 0.045 m was customized to each point cloud for segmenting unchanged and changed parts (see Fig. 2f and 2g).

Parallel to change detection, the point clouds were also used to create quantitative structural models (QSMs) of the trees (see Fig. 3a) by TreeQSM (Raumonen et al., 2013) in MATLAB (The MathWorks Inc., 2023) (see Fig. 3b). Some other QSM reconstructing programs like AdTree (Du et al., 2019) and AdQSM (Fan et al., 2020) build tree structures by the Dijkstra’s shortest path and the minimum spanning tree. They appeared to be sensitive to outliers in our dataset. Especially when they built detailed twigs at the branch’s high end, many shoots were invented by the algorithm due to outliers, not reflecting the actual sprouting pattern. In comparison to them, TreeQSM fits cylinders to point patches in defined sizes. This approach performs better in noise and outlier resistance than those methods using the Dijkstra’s shortest path. For each point cloud, we tested 18 configurations of different settings regarding the patch sizes for reconstructing the QSMs in TreeQSM. For each configuration, the reconstruction was repeated 15 times to reduce the impacts of pseudo-random numbers in this TreeQSM process. Finally, the QSM with minimum mean distances from points to trunk and branch cylinders was chosen as the model for the corresponding point cloud. It should be addressed again that each tree is represented with two point clouds, and accordingly two QSMs, showing their situations in 2022 and 2023 respectively. To further ensure a precise reconstruction, the outliers were pre-deleted through the statistical outlier removal tool (Rusu & Cousins, 2011).

**Figure 3.**
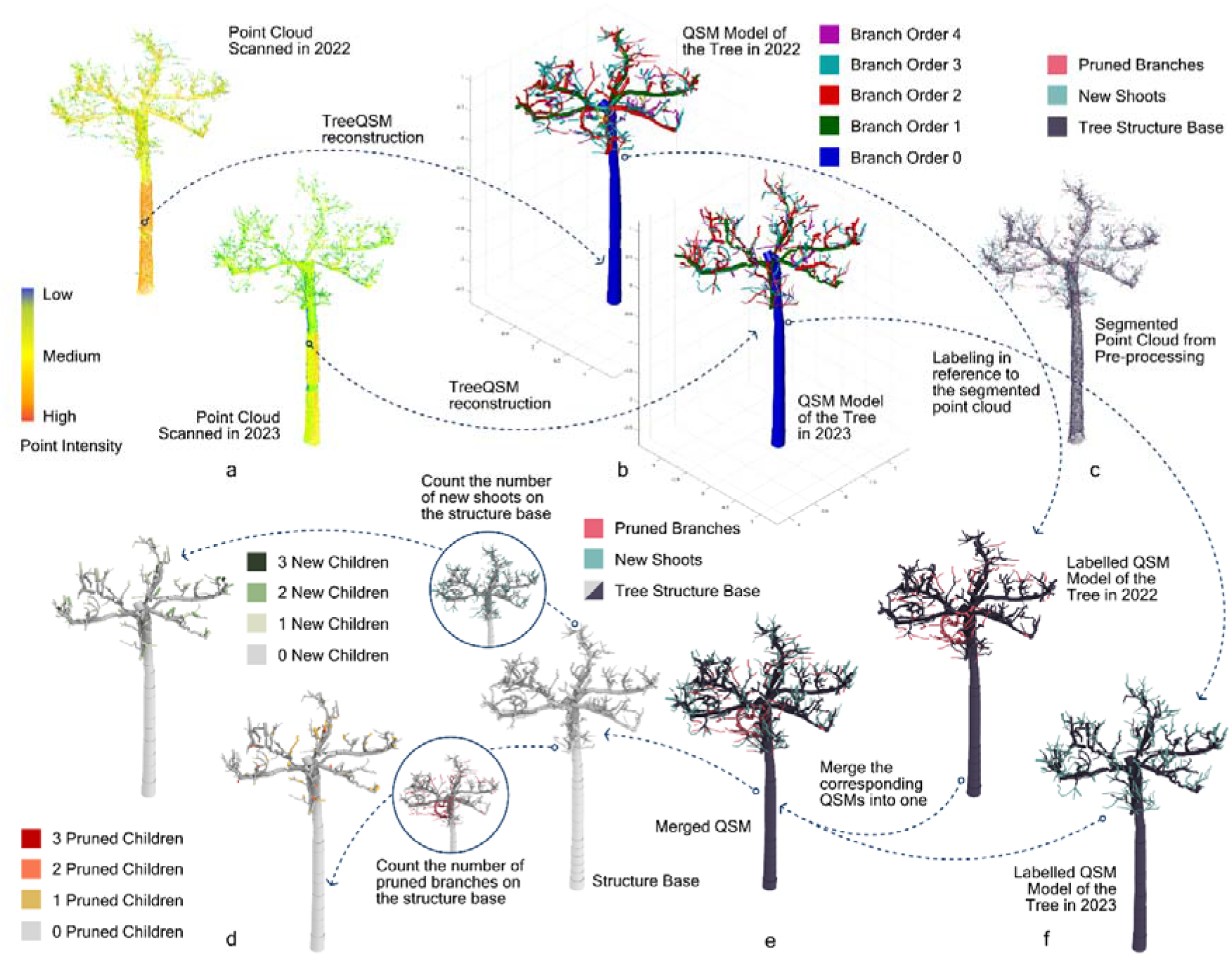
Overall procedure for labelling and reorganizing the dataset.

### 2.3 Labelling and reorganizing the dataset

In the pre-processing, the trimmed branches as well as the new shoots were detected in the point clouds while topological cylinders were generated with TreeQSM. The next step was to combine these two datasets. The individual cylinders of the QSMs must be labelled as to whether they are part of an unchanged branch (not considering the girth growth), a pruned branch or a new shoot. This was achieved by making use of a distance threshold between points of the cylindrical axis and their nearest neighbouring point of the segmented point clouds. For our data, we examined only the start and the end point of every cylinder. If the sum of their mean distances to their 10 nearest neighbours with the same label (i.e., trimmed branches) was below 100 mm, this cylinder was also labelled the same (see Fig. 3c 3f). To enhance the accuracy of the labelling, three more criteria were added based on practical rules when pruning these trees: for any cylinder labelled as part of either a new shoot or a pruned branch, its radius must be smaller than 20 mm (one year old shoots do not reach more than 20 mm in diameter for the trees at hand); for any cylinder labelled as part of a pruned branch, its branch hierarchical order must be larger than 1 (not the tree trunk and the primary branch); the label for trimmed branches and new shoots on one cylinder is passed on to all its children cylinders.

After labelling, the cylinders of different labels (unchanged branches, pruned branches and new shoots) are still separated in two QSMs regarding the same tree. There are no correspondences between these two QSMs as their reconstruction processes are independent. Therefore, cylinders of the trimmed branches in one QSM must be integrated into the other QSM that contains the main tree structure and the new shoots, or reversely, cylinders of new shoots must be integrated into the QSM with the trimmed branches. This is a tricky process. While the geometric data remain the same for every cylinder, its topological data regarding the ID of the cylinder, its parent cylinder and its child cylinder must be corrected, as well as the branch order and its position in the branch. Regarding whether to transfer cylinders of new shoots or pruned branches to the other QSM, considerations can be described as follows. The pruned branches, in general, could only be the same size or thicker than the new shoots. Consequently, cylinders of pruned branches have higher robustness in their position through cylinder fitting. As a result, the certainty for redefining their topological parent in another QSM based on their relative positions is supposed to be higher. So, for our dataset, the cylinders of pruned branches were picked out from their original QSM and integrated into the other QSM that has the new shoot cylinders (see Fig. 3e). Their new parent cylinders were redefined as those whose end points were located closest to their starting point. Based on this, the topological data for every single cylinder in the newly merged QSM were completely overwritten due to this change.

Finally, the total number of pruned branches and new shoots on every cylinder was counted (see Fig. 3d). This became the crucial attribute for the prediction models in the next step.

### 2.4 Prediction with various classification models

The dataset after all the processes described above contains 34,245 items, representing 28 table topped plane trees. Each item corresponds to one cylinder, which contains the following attributes: tree’s ID; cylinder’s ID; parent cylinder’s ID; child cylinder’s ID in the same branch; x-y-z coordinate of the cylinder start; a normalized 3d vector of the axial direction; branch’s ID; its sequence in the branch; branch order; cylinder length; cylinder radius; the number of pruned children and new children; the Boolean value if this cylinder is virtually added during QSM reconstruction; the Boolean value if this cylinder is pruned out.

The relationships between each two attributes (except for the IDs and Boolean values) are illustrated in appendix 2. For our research purpose, the sprout location and numbers are the label of new shoots on each cylinder. We tested classification models in machine learning for finding links between these topological and geometrical attributes and the predicting target. Among these target labels, 16,183 (47.3%) cylinders were labelled “-1” meaning that they are trimmed away. These cylinders are not feeding into machine learning models. 15,348 (44.8%) cylinders have no new shoot, thus labelled with “0”. 2,329 (6.8%) cylinders have one new shoot (labelled “1”). There are less cylinder samples, whose new shoot number is larger than “1”: 321 (0.94%) cylinders have 2 new shoots; 54 (0.16%) cylinders have 3 new shoots; 7 (0.02%) cylinders have 4 new shoots; 2 cylinders have 5 new shoots; only 1 cylinder has 6 new shoots on it. Due to the extreme rare samples with a high number of new shoots, we label those cylinders that have more than 4 shoots with new shoot number 4.

Owing to the limited volume of data we acquired, the majority of the items labelled with new shoot number from “0” to “4” must feed into machine learning models (16,558 items representing 26 trees). Nevertheless, we reserved 2 trees (1,504 items) as an evaluation dataset. This evaluation dataset was only used for validating the results (see section 3), not for training the model. The dataset for machine learning was further divided into a training set (13,246 items) and a testing set (3,312 items, with a test size of 0.2). The testing set prevented overfitting the models to the given data.

For getting a quick overview of the performances across a wide range of classification models in machine learning on the dataset, we used lazy predict (Pandala, 2023) to run scikit-learn (Pedregosa et al., 2011) to compare 25 common classification models with their default settings, including GaussianNB, NearestCentroid and LGBMClassifier. Besides, we tested a basic Artificial Neural Network model (ANN) built with Keras (Chollet, 2015). It consisted of two hidden layers with 64 and 128 nodes respectively (see Fig. 4 left). In addition, to examine a graph neural network (GNN) model, the dataset for each tree was processed to a graph (Salama, 2021), where every cylinder item was a node connected to its parent and children (the node connection for one tree is illustrated in Fig. 4 right). These graph data were fed into a GNN model named “baseline classifier” (see table 2.1) including 39,512 trainable params and 1,174 non-trainable params.

**Figure 4.**
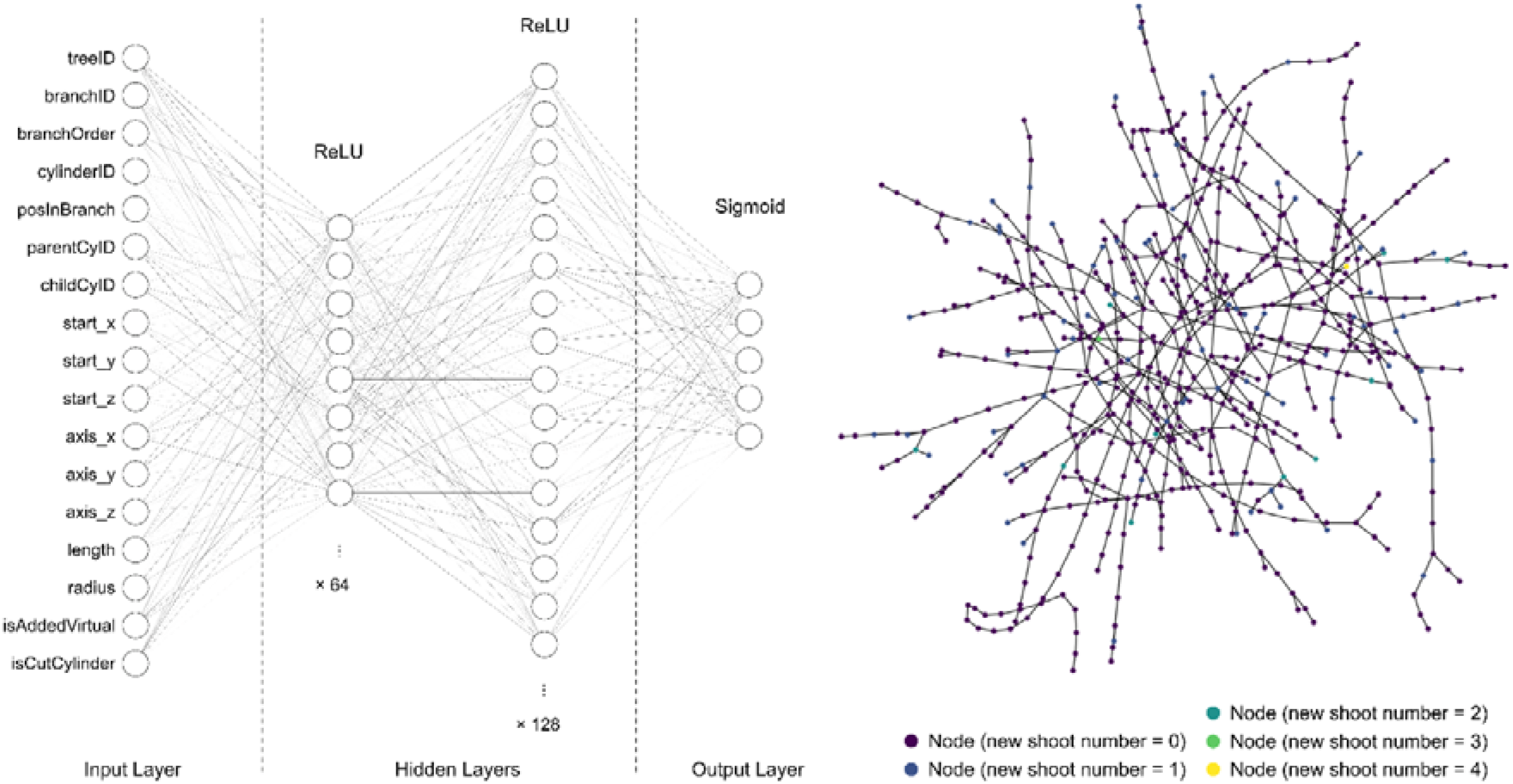
Architecture of the ANN (left) and Graph (right) of one tree used in our test.

**Table 2.1.** Architecture of the GNN model in our test.

| Layer (type) | Output Shape | Param | Connected to |
| --- | --- | --- | --- |
| input_features (InputLayer) | [(None, 11)] | 0 | [] |
| ffn_block1 (Sequential) | (None, 64) | 5228 | ['input_features[0][0]'] |
| ffn_block2 (Sequential) | (None, 64) | 8832 | ['ffn_block1[0][0]'] |
| skip_connection2 (Add) | (None, 64) | 0 | ['ffn_block1[0][0]', 'ffn_block2[0][0]'] |
| ffn_block3 (Sequential) | (None, 64) | 8832 | ['skip_connection2[0][0]'] |
| skip_connection3 (Add) | (None, 64) | 0 | ['skip_connection2[0][0]', 'ffn_block3[0][0]'] |
| ffn_block4 (Sequential) | (None, 64) | 8832 | ['skip_connection3[0][0]'] |
| skip_connection4 (Add) | (None, 64) | 0 | ['skip_connection3[0][0]', 'ffn_block4[0][0]'] |
| ffn_block5 (Sequential) | (None, 64) | 8832 | ['skip_connection4[0][0]'] |
| skip_connection5 (Add) | (None, 64) | 0 | ['skip_connection4[0][0]', 'ffn_block5[0][0]'] |
| logits (Dense) | (None, 2) | 130 | ['skip_connection5[0][0]'] |

We tested all these classification models in two manners of labelling: binary labels that only classify cylinders if they will or will not grow new shoots, and multiclass labels that classify cylinders based on the exact number of new shoots ranging between 0 to 4.

## 3. Results

The accuracy, balanced accuracy, and F1 Score (weighted average F1 score for multiclass labels) of the tested models in a default setting or with a basic architecture (see section 2.4) are listed in Fig. 5. Each scoring index ranges between 0 and 1. 1 is the best score, meaning that all the shoot labels are correctly predicted. On the contrary, 0 is the worst score, representing no correct prediction. In the figure, these models are shown in a descending order from the left to the right according to their total scores in classifying binary labels. Among the three sub-scores, accuracy reflects an overall rate of true predictions for all labels. Our datasets are imbalanced in terms of different label numbers. Therefore, balanced accuracy, which gives equal weights to the true prediction rates for each label, is also an important indicator in evaluating their performances. F1 score is another effective index for the imbalanced classifications but attaches more importance to true positives (predicting the cylinders with new shoots correctly) while it ignores the true negatives (predicting the cylinders with zero shoot correctly). Based on these benchmark scores, LGBMClassifier and GaussianNB have top scores for predictions with binary and multiclass labels respectively. The confusion matrix of the LGBMClassifier with binary labels in testing set is shown in table 3.1. The confusion matrix of the GaussianNB model with multiclass labels in testing set is shown in table 3.2.

**Figure 5.**
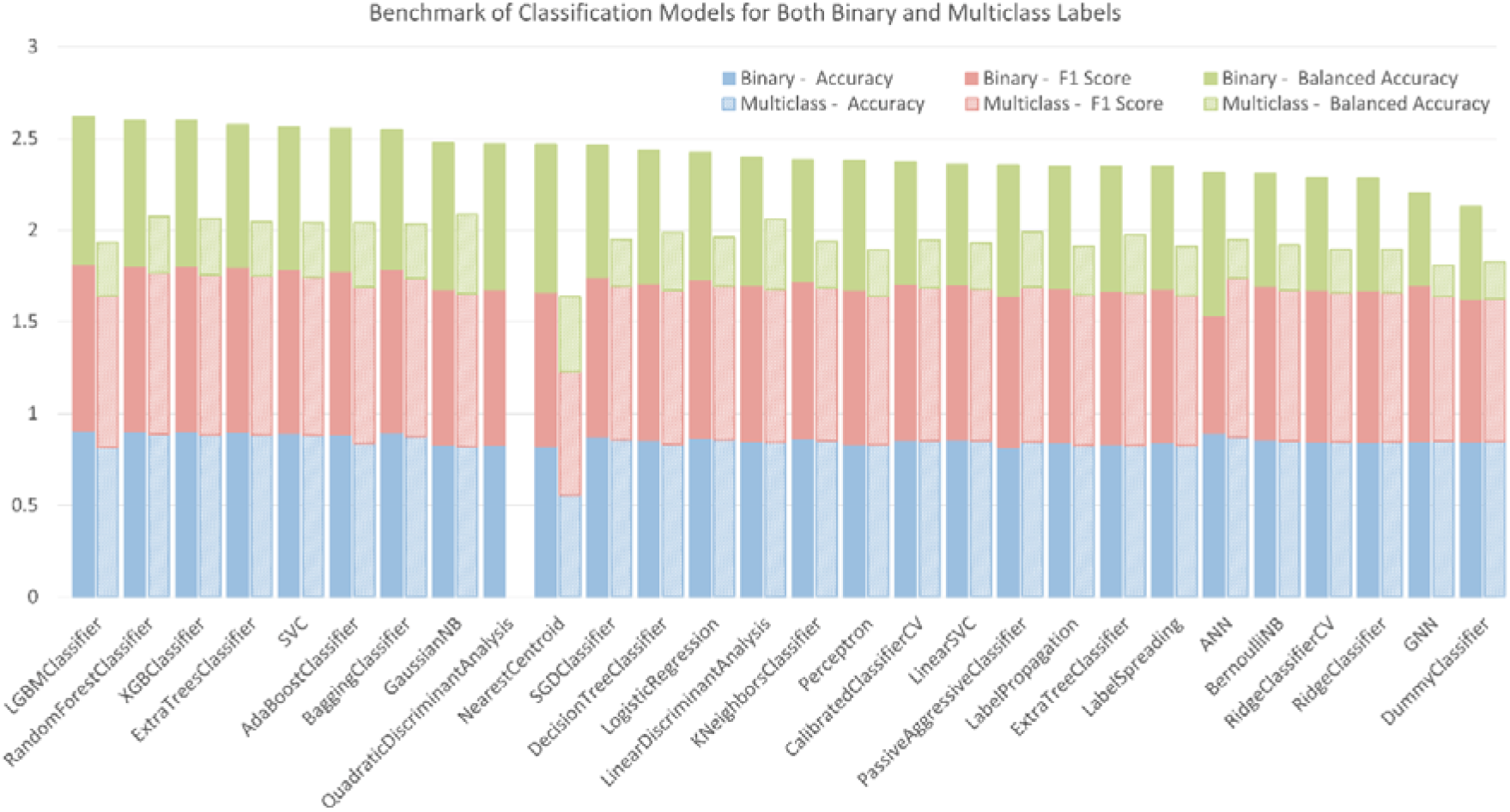
Benchmark of tested classification models for binary and multiclass labelling.

**Table 3.1.** Confusion Matrix of LGBMClassifier with Binary Labels in Testing Set.

|  | Negative / 0 | Positive / 1 |
| --- | --- | --- |
| Negative / 0 | 2883 (79.8%) | 170 (4.7%) |
| Positive / 1 | 209 (5.8%) | 351 (9.7%) |

**Table 3.2.** Confusion Matrix of GaussianNB Model with Multiclass Labels in Testing Set.

|  | 0 | 1 | 2 | 3 | 4 |
| --- | --- | --- | --- | --- | --- |
| 0 | 2458 (74.2%) | 309 (9.3%) | 33 (1.0%) | 0 (0%) | 1 (0%) |
| 1 | 153 (4.6%) | 258 (7.8%) | 24 (0.7%) | 0 (0%) | 3 (0.1%) |
| 2 | 29 (0.9%) | 27 (0.8%) | 5 (0.2%) | 0 (0%) | 0 (0%) |
| 3 | 4 (0.1%) | 4 (0.1%) | 0 (0%) | 0 (0%) | 1 (0%) |
| 4 | 2 (0.1%) | 1 (0%) | 0 (0%) | 0 (0%) | 0 (0%) |

To further validate these two models, we applied the trained LGBMClassifier model and GaussianNB model to the evaluation set with binary and multiclass labels respectively. The results of the evaluation are visually illustrated in supplementary material. Their accuracy, balanced accuracy and F1 Score on the validation set in comparison to the testing set is shown in table 3.3. The accuracy, balanced accuracy and weighted F1 score at the evaluation set (only 2 trees) have maximum around 10% difference to the scores on the benchmark.

**Table 3.3.** Scores of LGBMClassifier and GaussianNB Model in Evaluation Set.

|  | Scores in testing set |  |  | Scores in evaluation set |  |  |
| --- | --- | --- | --- | --- | --- | --- |
|  | Accuracy | Balanced accuracy | F1 Score | Accuracy | Balanced accuracy | F1 Score |
| LGBMClassifier with binary labels | 0.91 | 0.80 | 0.91 | 0.91 | 0.81 | 0.91 |
| GaussianNB Model with multiclass labels | 0.82 | 0.43 | 0.84 | 0.93 | 0.30 | 0.83 |

## 4. Discussion

In order to be able to meaningfully interpret and evaluate the results, it is first necessary to discuss the specific conditions of the dataset and resulting limitations.

The following factors may impact on the accuracy of the extracted geometrical data from the trees: 1) To prevent browsing the tree barks, protecting covers were installed at a height below 2 meters around the tree trunks. This might have caused the diameter measured at trunk cylinders to be slightly overestimated. However, we assume that this has no influence on the prediction model. 2) Minor swinging of the branches by wind during the LiDAR scanning might have caused outliers or might have led to overestimating the diameter of the smaller branches. Although the point clouds were denoised through SOR filters, this does not guarantee full deletion of these outliers and could then cause inexistent branches in the cylindrical models. 3) Aligning the same trees with different geometries in the two years is a nonstandard manual process so far, which can cause inconsistence in change detection and identification of parent cylinders. A possible alternative to detect these changes is comparing the occupancy grids (Hirt et al., 2021).

The total cylinder numbers for training the models were limited to 16,558 items representing 26 trees. The percentage of the negative label “0” makes up more than 92% of the total items, causing an unbalanced rate for the number of positive samples (less than 2500 items). Unfortunately, these are all available data from the nursery.

Most importantly, the collected dataset in two consecutive years reflects the growth of these trees under almost identical environmental conditions and pruning regime. More specifically, the temperature, water content in the soil, wind direction and speed as well as the time of pruning are all the same for these trees. This means that our method can predict the resprouting pattern of this kind of table-topped plane trees grown under the same conditions as in this study at current stage. In case of any changes in the factors mentioned above, it is unclear so far how accurate the prediction will be. For instance, the model may not predict the growth of the same trees in the following year, because horticultural experience shows that a change in the time of pruning of only one or two weeks can have a significant impact on the growth of new shoots, especially if there is also a change in weather conditions (e.g. heat or drought immediately after pruning).

For understanding whether those environmental factors could also be addressed in a prediction model in the same approach, these environmental data must be collected and coupled with a larger quantity of trees. This hints to an upcoming step of this study.

Finally, the current model is only the first step in understanding resprouting patterns after one specific artificial disturbance, namely pruning of table topped trees. Nonetheless we are optimistic that the approach has great potential for further development and application (see e.g., (Yazdi et al., 2023)). The application of such model is not limited to repeat what the gardeners can already do but go beyond knowledge boundaries regarding the resprouting strategy of trees after disturbances. This can hopefully be achieved through gathering a huge amount of data. By searching through this database, the “digital gardener” is likely to find evidence to support its predictions in a more complex context. For this far vision, an open-source and uniformed database about trees (Shu et al., 2022) is required.

## 5. Conclusion

Resprouting patterns are key in understanding regeneration of trees after natural and artificial disturbances. The interrelationships are very complex, involving among other things the primary status of hormones, the redistribution of resources, and timing issues. Until now, no single model can address all these factors in a physiological approach. However, gardeners and practitioners have been trained to prune trees based on their intuitive predictions since centuries. They are able to do so based on accumulated knowledge working with trees. In this study, we gave it a first try addressing the question if computational models, especially machine learning models could gain similar knowledge as practitioners from horticulture: what are the location and numbers of new shoots after pruning? Which model would achieve the best performance?

For this purpose, we scanned a group of annually pruned plane trees at a tree nursery with LiDAR. The detailed geometry and topology of the branches were extracted through quantitative tree models. The trimmed branches and new shoots were detected through comparison between the scans in two consecutive years and this information were finally labelled on a dataset for training multiple classification models.

We tested 25 common classification models in the field of machine learning with default settings. Additionally, 1 ANN model and 1 GNN model with most basic architectures were also tested. Among these models, except for two, all other models have an accuracy and a F1 score higher than 80%. For balanced accuracy, the average score of all the models was ca. 70% for binary labels; for multiclass labels, the average was 28.3%.

From the results, we can conclude that for the collected dataset, most of the models work well in telling the position of new shoots but are not accurate in telling the actual shoot numbers at the specific location. For the best scored models with binary labelling, the LGBMClassifier can predict the position of new shoots with an accuracy of 90.8% and a balanced accuracy of 80.3%. For predicting the exact number of the shoots, GaussianNB Model performs the best. The accuracy is 82.1% because most of the cylinders should have the shoot number 0. However, the balanced accuracy is reduced to 42.9%.

The limitation of the current model is definitely very specific to the studied site, environmental conditions, tree species and form, and the pruning time. In the next step, a larger amount of tree data is being collected in the city of Munich to analyse how this approach can be extended to a larger scope addressing maybe some of the environmental factors. In a further vision, a huge database of the “digital gardener” would push forward the knowledge boundaries in understanding resprouting strategies of trees facing natural and artificial disturbances.

## Supporting information

Supplemental raw data

## Acknowledgement

This study is funded by DFG-DACH-project under No. DFG-GZ: LU2505/2-1 AOBJ:683826 and cooperated with the DFG funded research groups under No. GRK 2678/1, AOBJ:68242, LU2505/2-1, PR 292/23-1 and RO 4283/2-1. Great credit to Bruns Pflanzen, who provided the table-topped plane trees as studying cases. This study would not have been possible without their dedication to scientific research and supports. Thanks to Luke Bohnhorst, who coordinated the use of RIEGL LiDAR scanner and the license of RISCAN Pro with us. Finally, the deep learning tutoring materials by Amir Ali and the graph Neural Networks tutoring materials by Khalid Salama were of great help to this study.

## Supplementary Materials

**Appendix 1.**
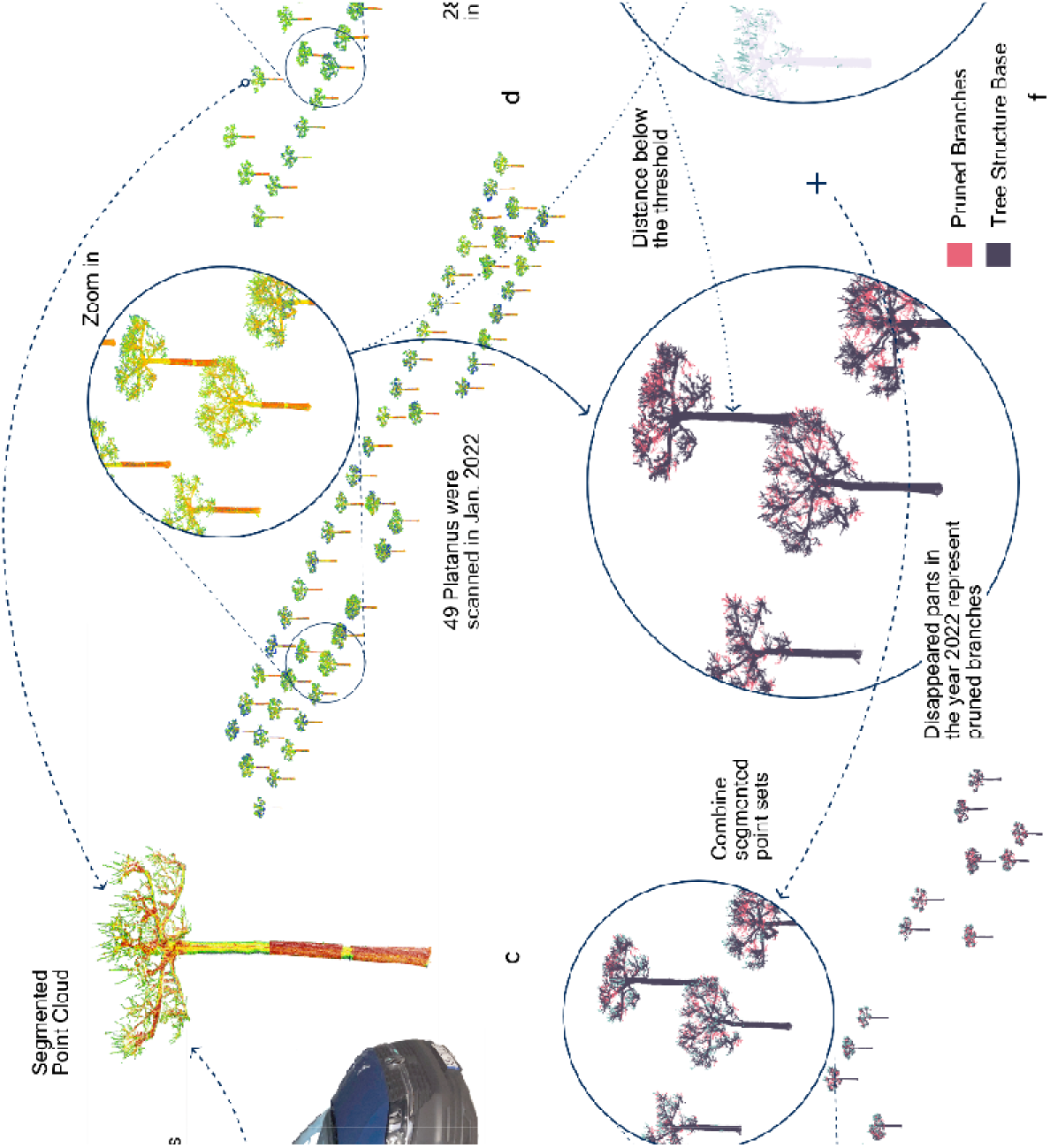
An overview of the workflow for point cloud acquiring and pre-processing

**Appendix 2.**
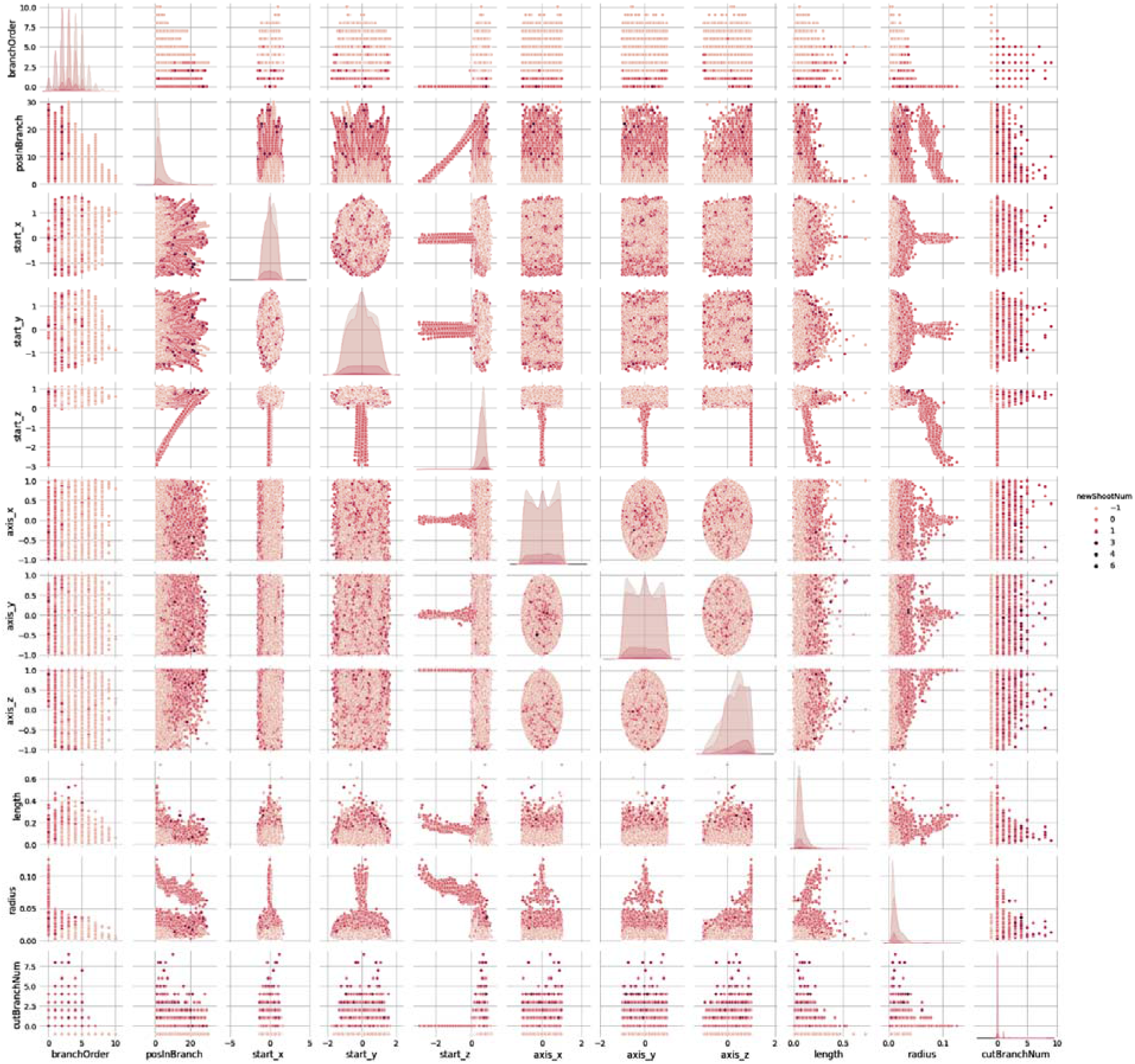
Correlations between each two attributes of the dataset. The number of new shoots is indicated with the colour, a darker colour represents a larger number of new shoots on the cylinder. The value “-1” is marked as pruned out cylinders.

**Appendix 3.**
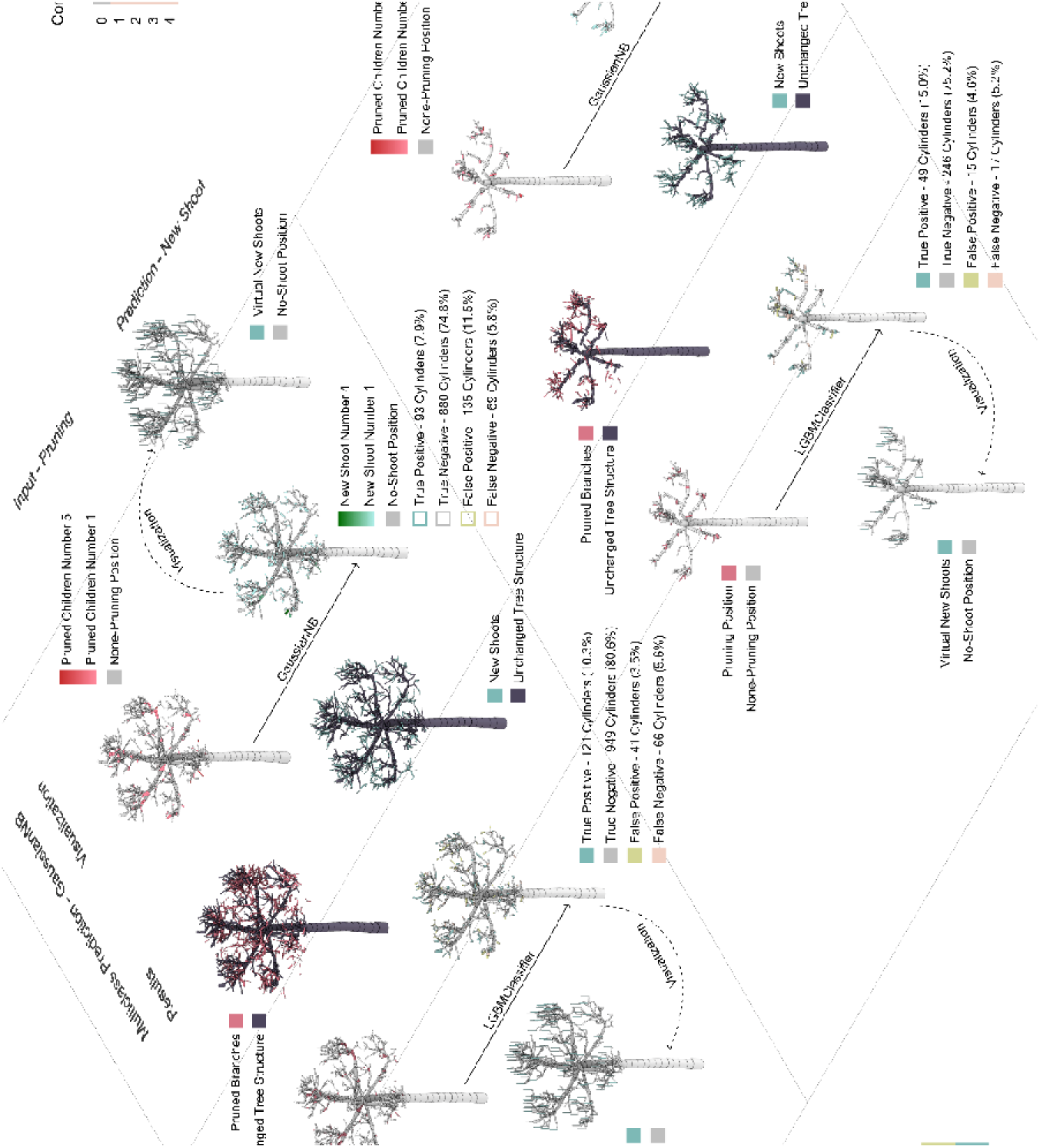
Visualizing the evaluation of the LGBMClassifier and GaussianNB models in predicting resprouting patterns. The ground truths are illustrated in the middle. The binary predictions using LGBMClassifier are drawn on the lower left side while the multiclass predictions using GaussianNB model are drawn on the upper right.

