## Supplementary figures and images for "Predicting resprouting of *Platanus* × *hispanica* following branch pruning by means of machine learning"

### Appendix 1

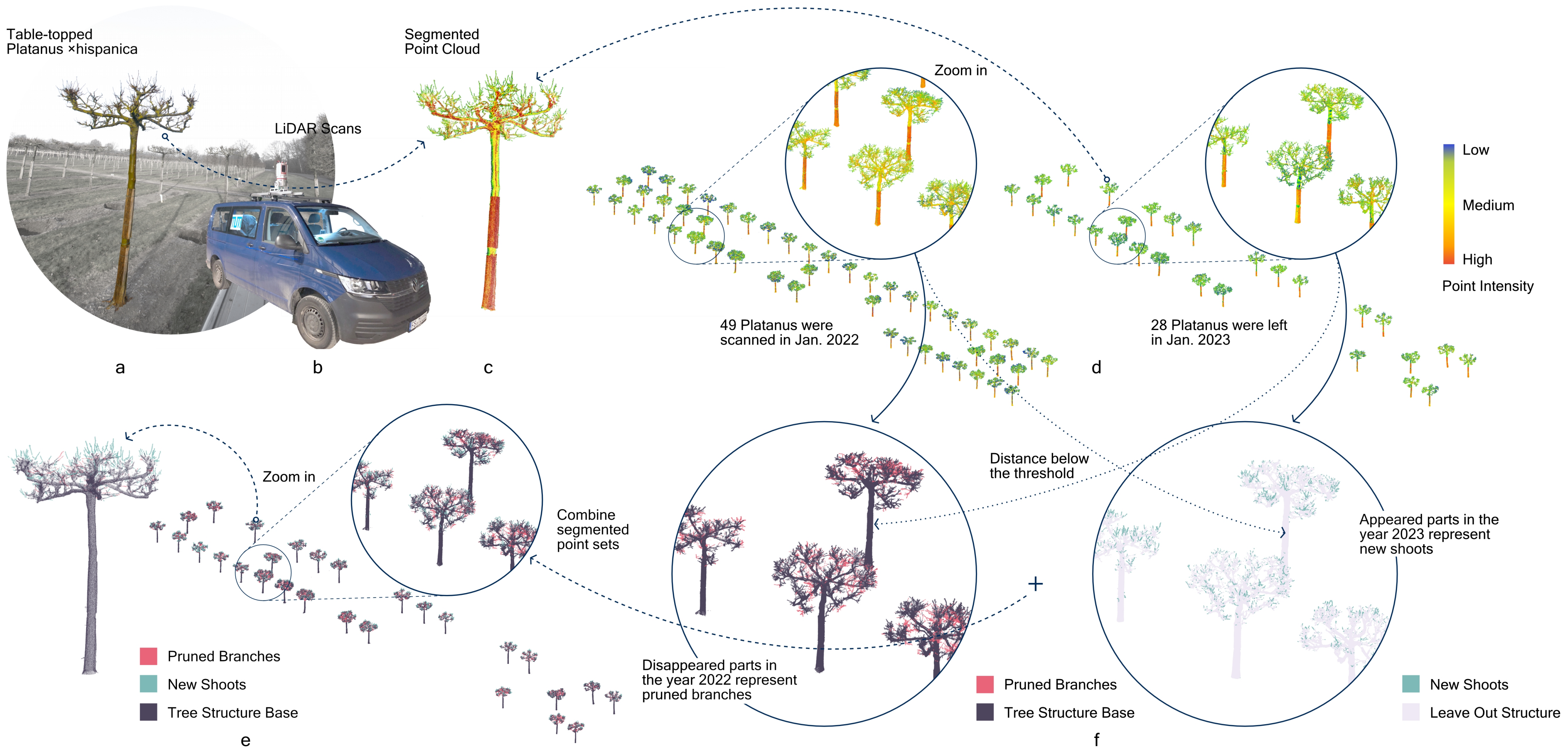

### Appendix 2

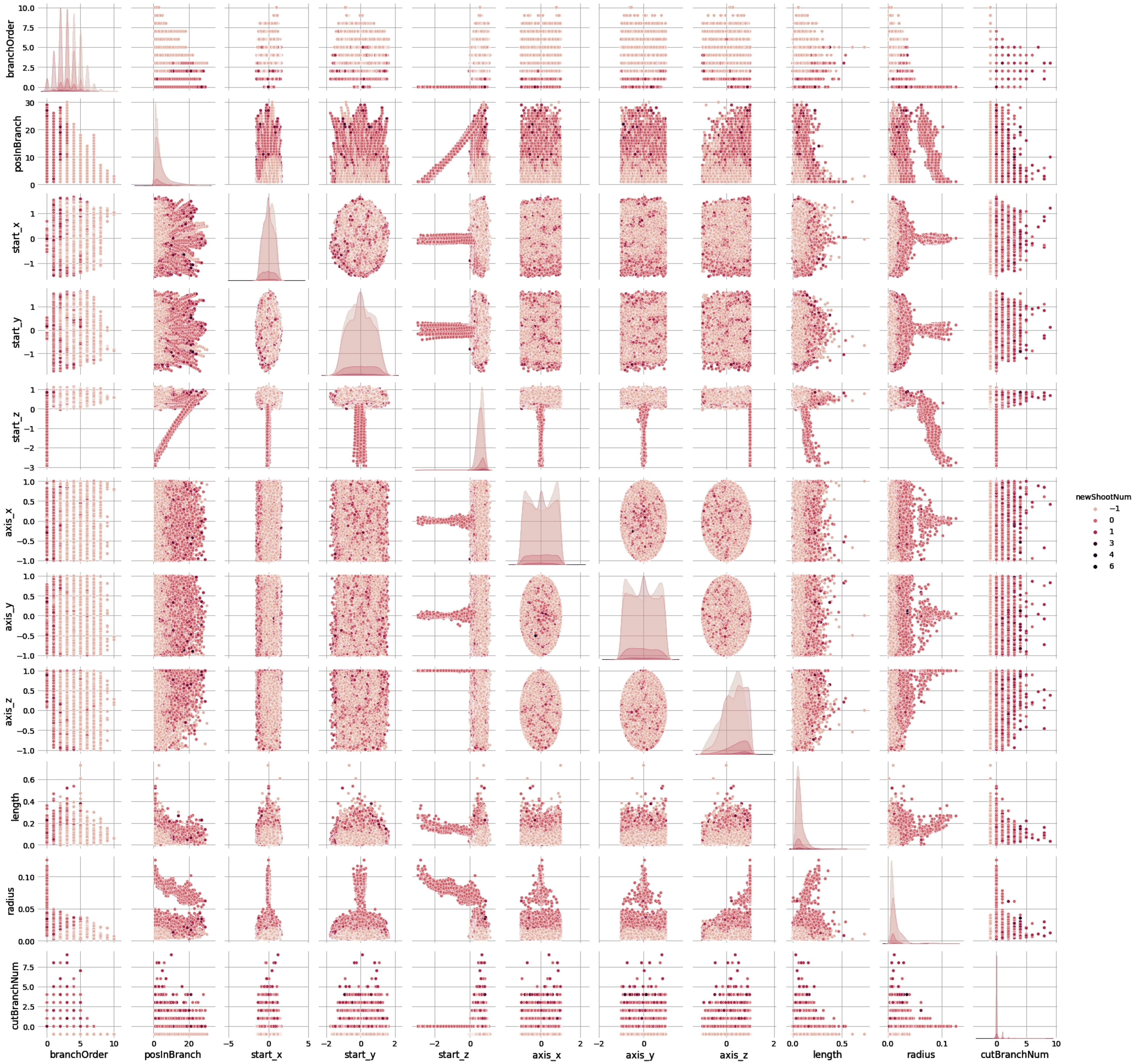
