## Appendix 3 for "Predicting resprouting of *Platanus* × *hispanica* following branch pruning by means of machine learning"

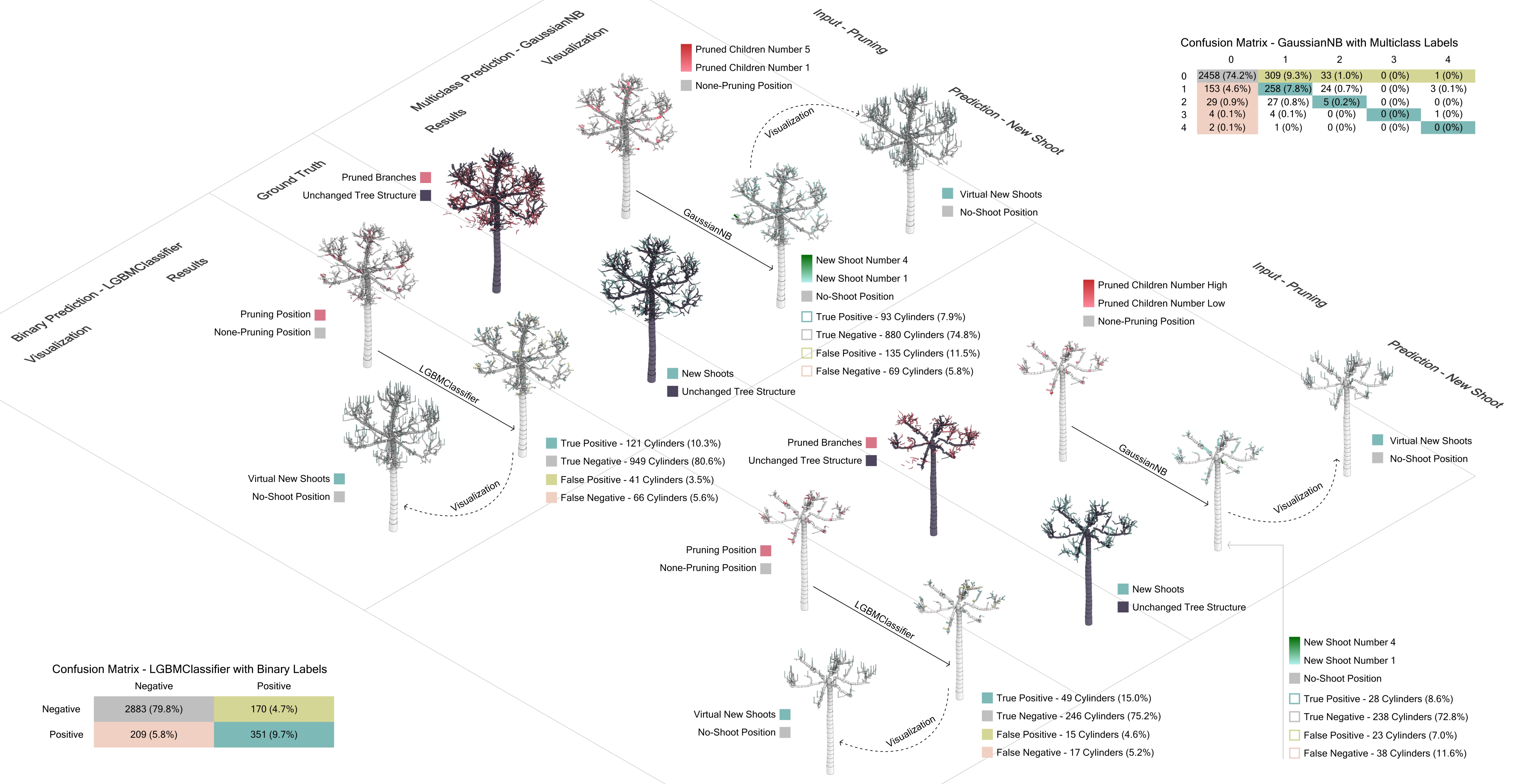


Confusion Matrix - GaussianNB with Multiclass Labels

| | 0 | 1 | 2 | 3 | 4 |
|---|---|---|---|---|---|
| 0 | 2458 (74.2%) | 309 (9.3%) | 33 (1.0%) | 0 (0%) | 1 (0%) |
| 1 | 153 (4.6%) | 258 (7.8%) | 24 (0.7%) | 0 (0%) | 3 (0.1%) |
| 2 | 29 (0.9%) | 27 (0.8%) | 5 (0.2%) | 0 (0%) | 0 (0%) |
| 3 | 4 (0.1%) | 4 (0.1%) | 0 (0%) | 0 (0%) | 1 (0%) |
| 4 | 2 (0.1%) | 1 (0%) | 0 (0%) | 0 (0%) | 0 (0%) |

Confusion Matrix - LGBMClassifier with Binary Labels

| | Negative | Positive |
|---|---|---|
| Negative | 2883 (79.8%) | 170 (4.7%) |
| Positive | 209 (5.8%) | 351 (9.7%) |
